# The Single-Cell Landscape of Peripheral and Tumor-infiltrating Immune Cells in HPV- HNSCC

**DOI:** 10.1101/2025.01.14.632928

**Authors:** Rômulo Gonçalves Agostinho Galvani, Adolfo Rojas, Bruno F. Matuck, Brittany T. Rupp, Nikhil Kumar, Khoa Huynh, Carlos Alberto Oliveira de Biagi Junior, Jinze Liu, Siddharth Sheth, Jelte Martinus Maria Krol, Vinicius Maracaja-Coutinho, Kevin Matthew Byrd, Patricia Severino

**Author notes:** Correspondence to: Kevin Matthew Byrd Patricia Severino. Equally contributed (Rômulo Gonçalves Agostinho Galvani, Adolfo Rojas).

## Abstract

Head and neck squamous cell carcinoma (HNSCC) is the sixth most common cancer worldwide. HPV-negative HNSCC, which arises in the upper airway mucosa, is particularly aggressive, with nearly half of patients succumbing to the disease within five years and limited response to immune checkpoint inhibitors compared to other cancers. There is a need to further explore the complex immune landscape in HPV-negative HNSCC to identify potential therapeutic targets. Here, we integrated two single-cell RNA sequencing datasets from 29 samples and nearly 300,000 immune cells to investigate immune cell dynamics across tumor progression and lymph node metastasis. Notable shifts toward adaptative immune cell populations were observed in the 14 distinct HNSCC-associated peripheral blood mononuclear (PBMCs) and 21 tumor-infiltrating immune cells (TICs) considering disease stages. All PBMCs and TICs revealed unique molecular signatures correlating with lymph node involvement; however, broadly, TICs increased ligand expression among effector cytokines, growth factors, and interferon-related genes. Pathway analysis comparing PBMCs and TICs further confirmed active cell signaling among Monocyte-Macrophage, Dendritic cell, Natural Killer (NK), and T cell populations. Receptor-ligand analysis revealed significant communication patterns shifts among TICs, between CD8+ T cells and NK cells, showing heightened immunosuppressive signaling that correlated with disease progression. In locally invasive HPV-negative HNSCC samples, highly multiplexed immunofluorescence assays highlighted peri-tumoral clustering of exhausted CD8+ T and NK cells, alongside their exclusion from intra-tumoral niches. These findings emphasize cytotoxic immune cells as valuable biomarkers and therapeutic targets, shedding light on the mechanisms by which the HNSCC sustainably evades immune responses.

## Introduction

Head and neck squamous cell carcinoma (HNSCC) is the most prevalent malignancy in the head and neck region, driven by a complex interplay of genetic and environmental factors. Risk factors associated with HNSCC in the oral cavity and larynx include tobacco use, alcohol consumption, environmental pollutants, and human papillomavirus (HPV) infection.^1–3^ These factors have led to the classification of HNSCC into two primary categories: HPV-associated (HPV-positive; HPV+) and HPV-unassociated (HPV-negative; HPV−) disease.

While HPV+ HNSCC generally has a more favorable prognosis, patients with HPV− HNSCC experience significantly poorer survival rates and face unique clinical challenges.^4^ HPV− HNSCC is marked by frequent mutations in key regulatory genes, including *TP53*, p16^INK4a^*(CDKN2A)*, and *CCND1*, with alterations in 60–80% of cases.^5^ These mutations drive tumorigenesis through mechanisms that are independent of HPV, in contrast to HPV+ HNSCC, where the viral oncogenes promote cancer development by inactivating key tumor suppressor proteins: E6 targets TP53 degradation, while E7 targets the retinoblastoma protein (RB). This disruption of tumor suppressor function is a hallmark of HPV+ cancers.^5–7^

Molecular distinctions extend to the tumor microenvironment (TME) and immune landscape, particularly in the behavior of tumor-infiltrating immune cells (TICs), which encompasses tumor-filtrating lymphocytes (TILs). HPV− HNSCC has been linked to a dysfunctional immune response, particularly involving CD8+ T cells, which are crucial for anti-tumor immunity.^8–10^

Lymph node (LN) metastasis, common in HNSCC, is associated with complex morphological adaptations in HPV− cases, including epithelial-to-mesenchymal transition, which results in subsequent tissue invasion influenced by immune evasion, altered cancer cell metabolism, and the upregulation of motility pathways, enabling tumor spread and immune resistance.^11–13^ Standard treatments for HNSCC, including surgical excision followed by chemotherapy or chemoradiotherapy, provide limited efficacy, particularly for advanced HPV− HNSCC.^14–16^

Despite advances in immunotherapies, such as immune checkpoint inhibitors (ICIs) that target the PD-1/PD-L1 pathway, survival rates remain poor, and therapeutic resistance, including immune cell exclusion from the tumor core, poses significant barriers.^17,18^

Pembrolizumab, an ICI approved for HNSCC, has demonstrated improved survival when combined with chemotherapy compared to cetuximab-chemotherapy regimens, yet outcomes remain suboptimal for metastatic HPV− HNSCC.^19,20^

Given these challenges, elucidating the immunological mechanisms within the TME, is critical to improving therapeutic outcomes for HPV− HNSCC. In this study, we focus on HPV− HNSCC to investigate immune cell dynamics across immune compartments (i.e., peripheral blood mononuclear cells versus lesion-associated TICs), nodal involvement and tumor stage. By integrating two previously published single-cell RNA sequencing (scRNA-seq) datasets,^21,22^ we analyzed a comprehensive cohort of 284,437 cells, comprising 177,191 TICs and 107,246 peripheral blood mononuclear cells (PBMCs) from 29 samples.

While we observed key differences in cell and molecular profiles in PBMCs and TICs. The differences between PBMCs and TICs/TILs became more pronounced as the disease progressed to nodal involvement or further stage, with the latter showing heightened immunosuppressive features within the tumor microenvironment.

Furthermore, we observed significant shifts in TIC subpopulation proportions as the disease advanced, with differential expressions of genes such as *KLRG1* and *FCRL6*, associated with T cell costimulation and the PD-1/PD-L1 pathway, respectively. All PBMCs and TICs revealed unique molecular signatures correlating with lymph node involvement; however, broadly, TICs increased ligand expression among effector cytokines, growth factors, and interferon-related genes. Pathway analysis comparing PBMCs and TICs further confirmed active cell signaling among Monocyte/Macrophages, Dendritic cell, Natural Killer (NK), and T cell populations. Receptor-ligand analysis revealed significant communication patterns shifts among TICs, between CD8+ T cells and NK cells, showing heightened immunosuppressive signaling that correlated with disease progression. In locally invasive HPV-negative HNSCC samples, highly multiplexed immunofluorescence spatial assays highlighted peri-tumoral clustering of exhausted CD8+ T and NK cells, alongside their exclusion from intra-tumoral niches. These insights lay the foundation for developing tailored immunotherapy strategies aimed at overcoming immune resistance mechanisms in this aggressive cancer subtype.

## Results

### Generating the First Integrated HNSCC Immune Cell Atlas

Due to factors such as patient diversity and cellular heterogeneity, HNSCC is a complex and difficult to study disease. Therefore, we set out to create an integrated HNSCC immune atlas using data from existing scRNAseq datasets.^21,22^ Each dataset was chosen because of its inclusion of both TICs and PBMCs samples, which represent the local and systemic immune environment within patients. For this study, we focus exclusively on HPV- HNSCC samples that were sorted for CD45+ immune cells from immunotherapy naïve patients. The overall demographics for this study included both sexes (62% males); primarily Caucasians (58.6%, others, unreported); a wide age range between 30- and 90-years-old with nearly half of the subjects between 50-69); and most of the tobacco and/or alcohol users included (69.0% and 51.7%, respectively; Table S1). The tumors, primarily T3/N0 tumors, were isolated from across the upper airways, with the majority isolated from the oral cavity. This diversity of demographic and disease data provided a foundation to explore immune cell diversity and states between peripheral and lesion-associated cell types as well as considering disease stage.

Before creating the integrated atlas, we first annotated and examined the cellular diversity across samples and pathological node status (N0/N+) to ensure no significant differences would hinder study integration. Using CellTypist,^23^ we identified and annotated all 14 PBMC and 21 TIC immune populations based on gene expression (Figure 1, Figure S1). PBMCs and TICs were visualized individually through UMAPs and stacked bar graphs (Figure S2, S3). Both studies demonstrated similar cellular frequencies and annotations in PBMC and TIC samples. Cells initially annotated as alveolar macrophages in GSE164690, characterized by high TREM2 expression and specificity for lung tissue, were reclassified as *TREM2+ Macrophages* for this study.

**Figure 1.**
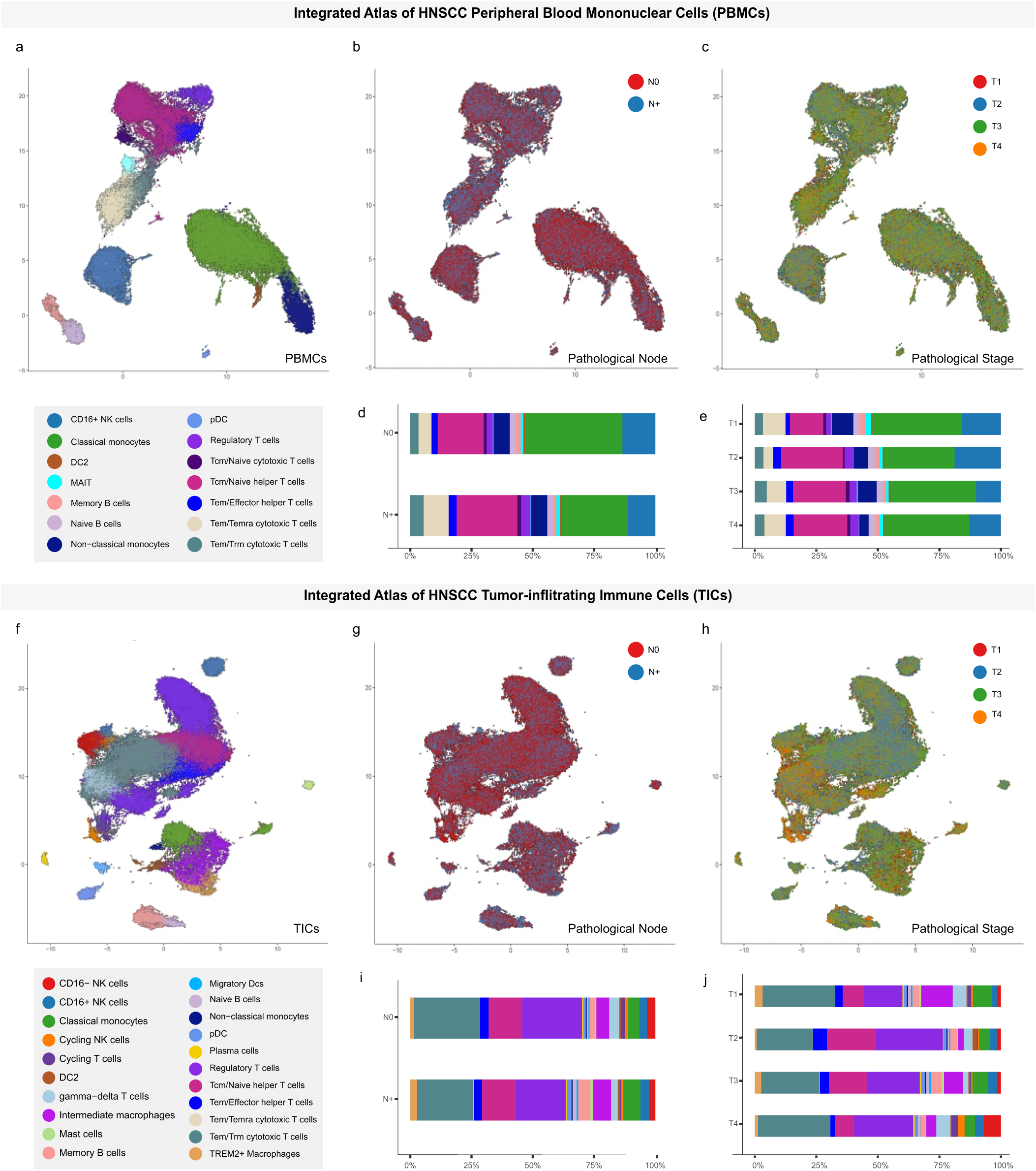
– Integrated Atlas of PBMCs and TICs in HNSCC samples. UMAPs colored by Cell Types (A), Pathological Node (B) and Pathologic Stage (C) in PBMCs. Bar graphs show average proportions of cell types across Pathological Node (D) and Pathologic Stage (E). UMAPs colored by Cell Types (F), Pathological Node (G) and Pathologic Stage (H) in TICs. Bar graphs show average proportions of cell types across Pathological Node (I) and Pathologic Stage (J). HNSCC – Head and Neck Squamous Cell Carcinoma; PBMC – Peripheral Blood Mononuclear Cells; NK – Natural Killer; pDC – Plasmacytoid Dendritic Cell; Tcm – T Central Memory; Tem – T Effector Memory; Trm – T Resident Memory; Temra – T Effector Memory expressing CD45RA; DC – Dendritic Cell; MAIT – Mucosal Associated Invariant T Cell; UMAP – Uniform Manifold Approximation and Projection; TIC – Tumor Infiltrating Cells;

After study specific integration, 29 samples from both studies meeting the above criteria were combined to create a HNSCC Immune Atlas using PBMC atlas consisting of 111,276 cells and one using TICs consisting of 167,565 immune cells (Figure 1). For the PBMCs and TICs atlas, cell signatures were extracted and UMAPs were generated according to cell types, pathological node (N0/N+ status) and pathological stage (T1-T4). The integrated PBMC datasets varied in cellular proportions across nodal involvement and pathological stage. N+ samples displayed higher proportions of Tem/Temra cytotoxic T cells compared to N0 samples, while T3 and T4 samples displayed lower proportions of CD16+ NK cells compared to lower stage samples (Figure S4A and S4B, respectively). Differences in cell proportions were also observed across node involvement within TICs. Specifically, N+ samples showed lower proportions of Tem/Trm cytotoxic, regulatory T cells (Tregs) and gamma-delta T cells, (Figure S4C) compared to N0 samples. When evaluating cell populations relative to T status, we did not find any differences (Figure S4D). In both peripheral and centralized compartments, these integrated datasets reveal some distinct immune cell shifts comparing nodal involvement or pathological stage in both TICs and PBMCs. However, the altered cell proportions comparing nodal involvement or pathological stage found in the PBMC samples were different from those found in the TICs, suggesting that the changes found in the latter may be the result of the tumor microenvironment or the tumor itself.

### Single-cell Signatures of Nodal Metastasis and Tumor Stage

The underlying biological cause of cancer metastasis in specific patients is elusive but hypothesized to be attributed to various signaling from the TME and circulating immune cell populations.^24^ While our analyses observed some differences in immune cell enrichment, we next wanted to explore molecular signatures of nodal involvement and pathologic stage (Figure 2, Figure S5). Broadly, major differences in gene expression signature were observed between N+ and N0 samples and T3/T4 versus T1/T2 stage, visualized using jitter plots (Table S2). Analyzing the TICs, the main DEGs upregulated in N+ when compared to N0 are genes related to the response to interferon, the most frequent being *IFIT3, IFIT2, IFIT1* and *IFITM1* expressed mainly by cells of the myeloid lineage such as Intermediate Macrophages and Classic Monocytes.^25,26^ In contrast, the genes mostly upregulated in the lymphoid population such as, but not limited to, Cycling NK, CD16-NK cells, Gamma-Delta T cells, Tem/Trm Cytotoxic T cells and Tregs are genes associated with immunomodulation such *as IL7R, KLRG1, KLRB1* and *IL12RB2*.^27–31^ In addition to genes associated with the response to interferon and immunomodulation, genes related to angiogenesis such as *VEGFA* and *FN1*, in DC2 and TREM2+ Macrophages, were found (Figure 2A).^25,26,32–35^

**Figure 2.**
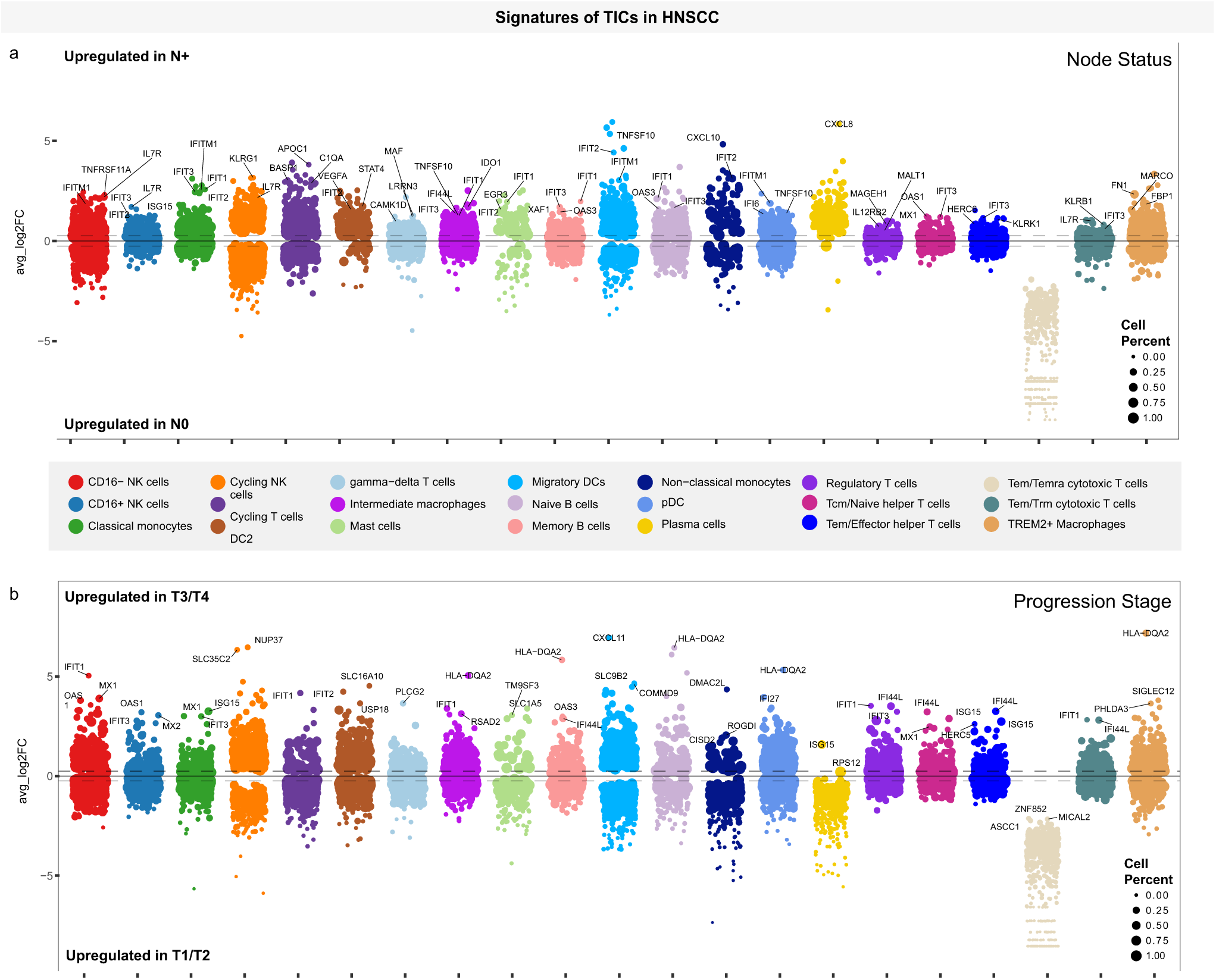
**- Differentially expressed genes at the single cell level in TICs**. Jitterplots indicating differentially expressed genes between N status (A) and Pathological Stages (B). The size of each dot represents the percentage of cells expressing the gene. Cell types have been highlighted by color. Genes were ranked based on the average Log2FC. Highlighted genes are those that showed the highest differential expression with known immunological function among the 20 genes with highest Log2FC in each cell annotation. The black line represents Log2FC of zero. The dotted line represents the Log2HR threshold of 0.25. HNSCC – Head and Neck Squamous Cell Carcinoma; NK – Natural Killer; pDC – Plasmacytoid Dendritic Cell; Tcm – T Central Memory; Tem – T Effector Memory; Trm – T Resident Memory; Temra – T Effector Memory expressing CD45RA; DC – Dendritic Cell; MAIT – Mucosal Associated Invariant T Cell.

Regarding the pathological stage, we again found that most of the upregulated DEGs in cells from individuals with tumors at a higher stage of progression (T3 or T4) are related to the response to interferon, with the most frequent being *IFIT1, IFI44, IFI44L* and *IFIT3,* however not mainly in cells of myeloid origin, but in lymphoid cells such as CD16- NK cells, CD16+ NK cells, Cycling T cells, Tregs, Helper T cells, Memory B cells and Plasma cells.^25,26^ In antigen-presenting cells such as Intermediate Macrophages, Naive B cells, pDC and TREM2+ Macrophages, overexpression of *HLA-DQA2* was found, a gene that encodes the low polymorphic alpha allele of HLA-DQ incapable of dimerizing with HLA-DQB1 (Figure 2B).^36,37^ However, when we compared the PBMC samples of individuals in relation to nodal involvement or pathological tumor stage, interferon response genes did not appear among the DEGs in the cell populations studied, with the exception of N+ Non-Classical Monocytes that overexpress *IFIT3*, *IFI44* and *IFI44L* (Figure S5). Furthermore, we did not observe a molecular signature pattern inherent to the cell type between PBMCs and TICs, suggesting that there is a modulation induced by the TME or the tumor itself that could be producing Interferons.

### Gene Set Enrichment Analysis of Immune Cells in HPV- HNSCC

To assess whether genes differentially expressed between the N+ and N0 groups influence key biological pathways, we conducted a Gene Set Enrichment Analysis (GSEA) using the MSigDB Hallmark library,^38,39^ which contains 50 pathway collections relevant to different cell types. As expected, differences in GSEA were observed between N+ and N0 samples and T3/T4 versus T1/T2 stage, visualized using bubble plots (Table S3 - Figure 3). Regarding N status, in the TICs atlas, 112 pathways were found to be enriched. Of these, 61 pathways were positively enriched in cells from the N+ group, with DC2 cells showing the highest enrichment with 18 pathway sets, followed by migratory DCs with 10. In contrast, 51 pathways were negatively enriched in the N+ group, with memory B cells having the most negative enrichments (nine pathways), followed by Tem/Effector Helper T cells with eight downregulated pathways. In PBMCs atlas, we found only 27 enriched pathways. Among these, six pathways were positively enriched in cells from the N+ group, with memory B cells showing three enriched pathway groups. On the other hand, 21 pathways were negatively enriched in cells from the N+ group, with MAIT cells having five negatively enriched pathway clusters. This analysis suggests that due proximity, the tumor resident immune cells are more modulated by the tumor than PBMCs, emphasizing the effect of the tumor on the surrounding environment, a concept being further defined as structural immunity in overall and even oral mucosal immunity research (Figure 3A).^40–42^

**Figure 3.**
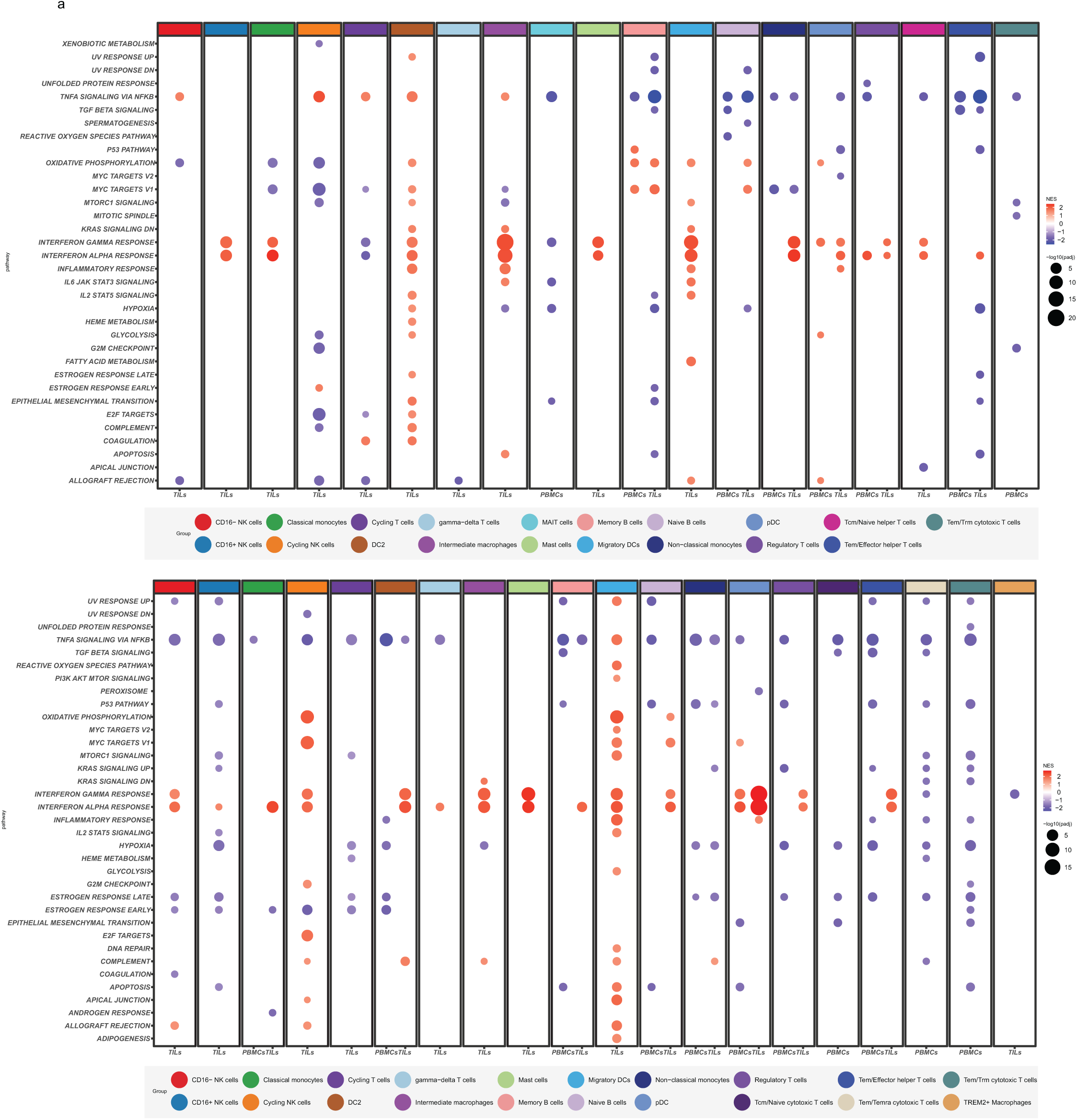
**- Gene Set Enrichment Analysis (GSEA)**. The terms were determined using the MSigDB_Hallmark library. A GSEA analysis was conducted per cell type, comparing N+/N0 cells (A) or T3+T4/T1+T2 (B) and considering all DEGs. The graphical representation illustrates the enrichments of cell types present in TICs from the integrated data sets of GSE139324 and GSE164690. The size of the points in the figure represents -Log padj value of each enrichment in that specific cell type and the color represents Normalized Enrichment Score (NES) from lowest (blue) to highest (red). NK – Natural Killer; pDC – Plasmacytoid Dendritic Cell; Tcm – T Central Memory; Tem – T Effector Memory; Trm – T Resident Memory; Temra – T Effector Memory expressing CD45RA; DC – Dendritic Cell; MAIT – Mucosal Associated Invariant T Cell.

Among the gene sets evaluated, three of them were those that showed enrichment across a wide variety of cell populations. TNFα signaling via NF-κB was enriched in 18 populations, 11 cell populations in the TIC atlas and seven tin the PBMC atlas. Of the TICs, five cell populations showed positive enrichment and six showed negative enrichment. In PBMCs, this gene set was negative. Two of the most prevalent pathways were Interferon Gamma and Alpha Responses (IFNγ and IFNα). IFNγ was observed in 11 TIC immune cell populations and two PBMC immune cell populations, while IFNα was enriched 14 times, 13 in TICs and once in PBMCs Regulatory T cells.

When comparing higher progression stage (T3-T4) to lower stage, 56 pathways were positively enriched samples in TICs sample, with Migratory DC being the most enriched with 19 pathways followed by Cycling NK with 9 enriched pathways. On the other hand, 37 pathways were negatively enriched in TIC T3-T4 group, with 10 negative enrichments in CD16+ NK cells. In the PBMC Atlas, only 3 pathways were positively enriched in T3-T4, all in pDC. In contrast, 67 pathways were negatively enriched, with Tem/Trm Cytotoxic T cells with 14 pathways and Tem/Temra Cytotoxic T cells with 13 pathways (Figure 3B). This analysis, once again, suggests tumor-proximal modulation is different from the modulation observed systemically. Among these pathways, the most enriched were Interferon Alpha Response, positively enriched in 14 cell populations in TICs and in only 1 population in PBMCs, Interferon Gamma Response, positively enriched in 10 cell populations in TICs and only 1 in PBMCs and TNF-α signaling via NFκB, with negative enrichments in 8 cell populations in TICs and 11 in PBMCs. In combination with the DEG data, this analysis demonstrates that interferon pathways are highly activated in TIC, again emphasizing a modulation of the immune system dependent on interferon in TME.

### Receptor-Ligand Networks in TICs

To investigate the communication dynamics within the TME, we focused on the TICs and analyzed the receptor-ligand signaling and predicted interaction networks between cell populations *CellChat*.^43^ The cell populations with the highest number of outgoing signals in the N0 group were Intermediate Macrophages and TREM2+ Macrophages (Figure 4A). A similar pattern was observed in the N+ group, where Intermediate Macrophages and TREM2+ Macrophages again showed the highest number of outgoing interactions (Figure 4B). For incoming signals, the same two populations, TREM2+ Macrophages and Intermediate Macrophages, exhibited the highest number of interactions in both N0 and N+. However, the most significant change in incoming interactions was observed in Temra/Tem Cytotoxic T cells, which showed an increase of over 70% compared to N0, while Cycling NK cells exhibited a reduction of almost 30% (Figure 4B).

**Figure 4.**
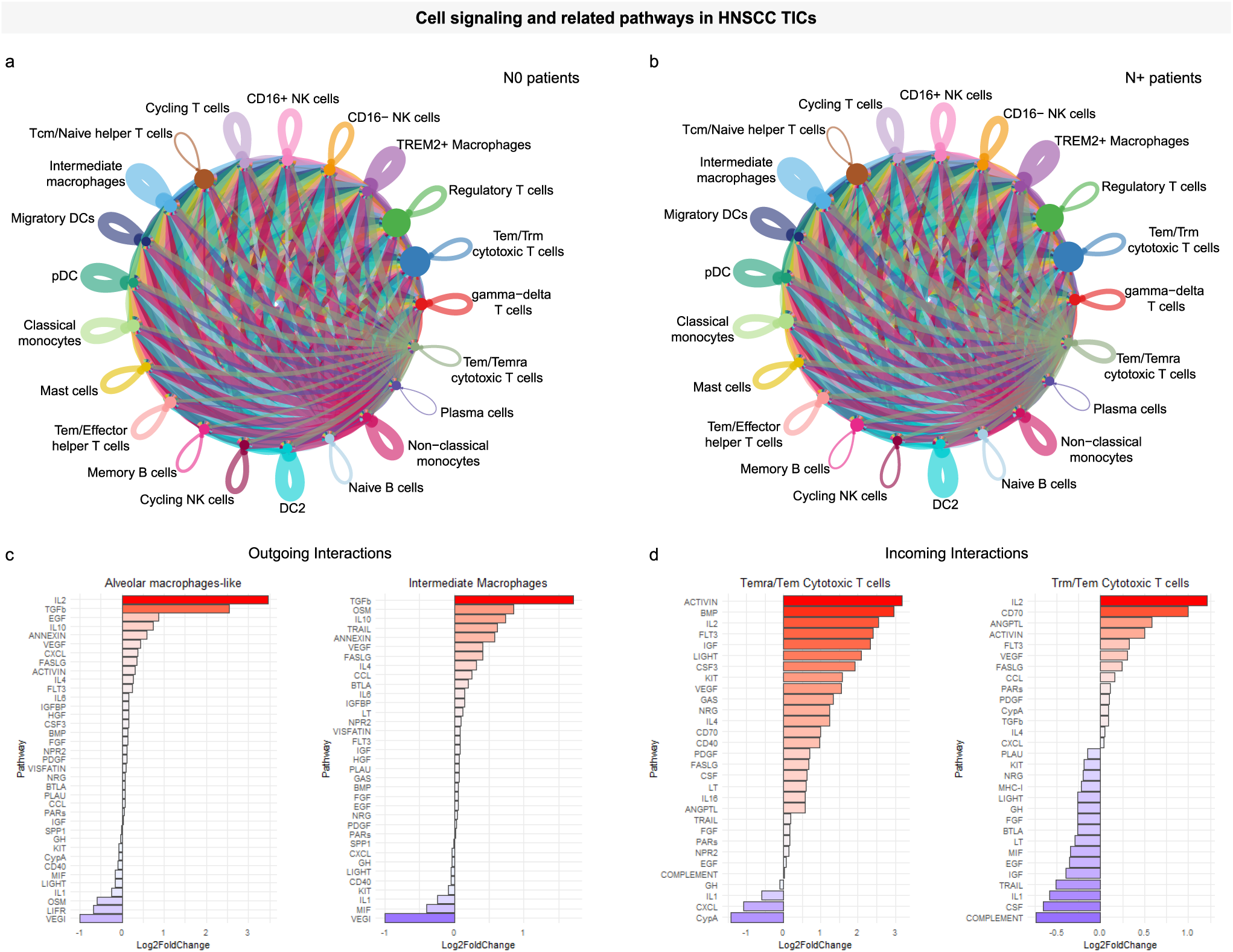
Cell-cell communication in TIC. Circular plots illustrating cellular communication among major cell types, representing the numbers of interactions between these cells in TIC from N0 (A) of N+ (B). Bar plots representing the Log2 Fold Change (N+ over N0) in outgoing interactions from Intermediate Macrophages and Alveolar/TREM2+ Macrophages (C) and incoming interactions from Trm/Tem and Temra/Tem Cytotoxic T cells (D). HNSCC – Head and Neck Squamous Cell Carcinoma; NK – Natural Killer; pDC – Plasmacytoid Dendritic Cell; Tcm – T Central Memory; Tem – T Effector Memory; Trm – T Resident Memory; Temra – T Effector Memory expressing CD45RA; DC – Dendritic Cell; MAIT – Mucosal Associated Invariant T Cell; UMAP – Uniform Manifold Approximation and Projection; TIC – Tumor Infiltrating Cells;

Macrophages are known to interact with T lymphocytes through antigen presentation but also have immunomodulatory roles.^44^ Given the increase in outgoing signals from macrophages and the rise in incoming interactions in Temra/Tem Cytotoxic T cells, we assessed which signaling pathways underwent the most notable changes. We focused on outgoing signals from TREM2+ Macrophages and Intermediate Macrophages, as well as incoming signals received by Temra/Tem Cytotoxic T cells and Trm/Tem Cytotoxic T cells. Our evaluation of the pathways related to outgoing signals from Intermediate Macrophages and TREM2+ Macrophages (Table S4) showed that the Transforming Growth Factor Beta (TGF-β) and IL-10 pathways demonstrated the largest signal increase in the N+ group compared to N0 (Figure 4C). For incoming signals in cytotoxic T lymphocytes, pathways associated with IL-2 and CD70, both linked to T lymphocyte activation, were elevated in Temra/Tem Cytotoxic T Cells and Trm/Tem Cytotoxic T Cells in N+ compared to N0 (Figure 4D). These data point to an increase in interactions with immunosuppressive potential from macrophages, and in interactions related to activation in cytotoxic T lymphocyte populations.

### Single-cell Trajectories of Cytotoxic Immune Cells

To investigate the roles of cytotoxic innate and adaptive immune cells in tumor progression, we analyzed the expression of key genes that regulate the activity and modulation of NK cells and cytotoxic T lymphocytes across different nodal and pathological stages. This was done through pseudotime and gene trajectory analysis. For NK cells (including CD16+, CD16-, and cycling NK cells), we evaluated the expression of genes related with NK cell activation or inhibition roles: *AREG, KLRG1, TCF7, GZMA*, and *KIR2DL4*. While NK cells from N0 individuals were evenly distributed throughout the pseudotime progression, those from N+ individuals were predominantly observed in later stages, suggesting a progression from N0 to N+ for NK cells along the pseudotime (Figure 5A-E). Notably, we observed an increased expression of immunomodulatory genes *AREG* and *KLRG1* over pseudotime, with higher levels in NK cells from N+ individuals shown by the line representing the trend. Similarly, *TCF7*, which is crucial for NK cell maintenance, showed greater expression in NK cells from N+ individuals. However, when analyzing *GZMA* and *KIR2DL4*, genes associated with NK cell cytotoxicity and activation, we found the opposite trend. A reduction in expression as pseudotime progressed, with lower levels of these genes in NK cells from N+ individuals. These findings suggest a shift toward an immunomodulatory role for NK cells in N+ individuals, with diminished cytotoxic activity compared to those from N0 individuals.

**Figure 5.**
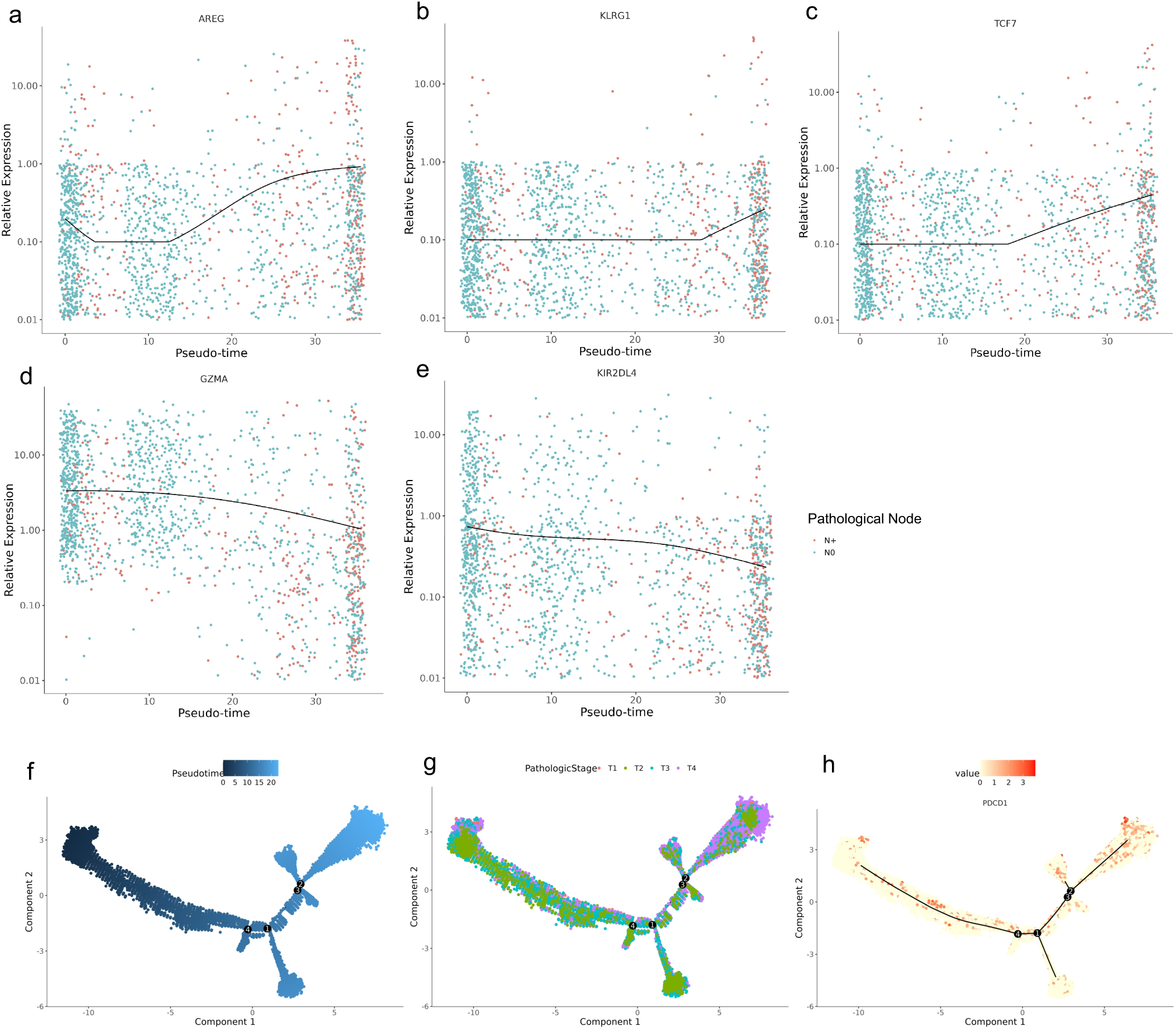
Gene trajectories in TICs. Jitterplots with average line illustrating relative expression of *AREG* (A), *KLRG1* (B), *TCF7* (C), *GZMA* (D) and *KIR2DL4* (E) in NK cells. Blue dots represent cells from N0, and red dots represent cells from N+. PCA representing gene trajectories of *CHUK, HLA-DRB5, HLA-DQA1, HLA-DQB1, CD3D, HLA-DRA, NFKBIA, HLA-DRB1, TRAC, UBE2N* in Cytotoxic T cells over pseudotime (F), Pathologic Stage (G) and *PCDC1* expression (H). PCA – Principal Component Analysis.

For cytotoxic T lymphocytes (including Temra/Tem and Trm/Tem cytotoxic T cells), we analyzed genes related to TCR downstream signaling: *CHUK, HLA-DRB5, HLA-DQA1, HLA- DQB1, CD3D, HLA-DRA, NFKBIA, HLA-DRB1, TRAC*, and *UBE2N*. The trajectory analysis revealed a distinct separation in pseudotime, where one trajectory, occurring later in pseudotime, contained more cells from individuals with T4 pathological stage disease (Figure 5F-H). Additionally, when analyzing the expression of *PDCD1* (PD-1), a critical marker of CD8 T cell exhaustion, we found that most cells expressing this marker were located along the same trajectory as the T4-stage cells. This suggests that as cytotoxic T cells progress toward the T4 stage, they adopt an exhausted phenotype, marked by high PD-1 expression. Together, these data highlight a shift in NK cells toward an immunosuppressive role in N+ individuals, alongside a trajectory toward T cell exhaustion in cytotoxic T lymphocytes, particularly in patients with advanced disease. This shift likely contributes to the diminished cytotoxic response observed in the tumor microenvironment of advanced-stage HNSCC.

### Comparison of results across scRNAseq and bulk RNAseq datasets

To both validate and ascertain the clinical relevance of these findings, we attempted to explore another existing dataset, we used the data deposited in The Cancer Genome Atlas Head-Neck Squamous Cell Carcinoma (TCGA-HNSC) project. To make the cohort comparable, we selected only samples from HPV-negative individuals, Caucasians, and with tumors in the same oral cavity sites as our study (Table S1) Using their bulk RNAseq data, we first assessed whether tumor stage (T4 x T1), nodal involvement or any of the genes examined in Figure 5 significantly increased. We did not observe statistical differences in the gene expression levels of the AREG, KLRG1, GZMA and KIR2DL4 genes when comparing the N0 and N+ samples, in the same way that we did not see any difference between the survival curves of both groups, although a better, non-statistically significant survival was observed in the N0 group (Figure S6A). Regarding the PDCD1 gene, when we compared its expression level between T1 and T4 individuals, no difference was observed, as in the survival curve, although again there was a trend towards better survival in individuals in the T1 group (Figure S6B).These results using bulk RNA reflect the lack of resolution of this approach, as compared with scRNAseq, for the identification of cell population-specific markers, suggesting the need for more in deep single and spatial analysis. We therefore attempted to validate our Single Cell RNAseq findings in Spatial Analysis by trying not only to validate the data, but to locate the tissue where these cells would be found in the tumor environment.

### Understanding the Spatial Enrichment of Cytotoxic Cell States in Situ

We observed single cell heterogeneity in our cytotoxic immune cells within the TICs, which we hypothesized would be connected to the spatial position of the immune cell within the TME. This is critical because studies have found prognostic potential of individual immune types cells when considering their position at the border of the tumor (peri-tumoral) versus within the tumor (intra-tumoral).^45^ To further investigate the role of cytotoxic immune cells in HPV-negative HNSCC, we conducted a focused multiplexed immunofluorescence (Multi-IF) experiment using the Phenocycler-Fusion 2.0 (Akoya Biosciences) to profile specific cytotoxic immune cell states within the tumor microenvironment. We utilized tissues from a subset of subjects enrolled in a clinical trial (NCT0352942 with locally invasive HPV-negative HNSCC with metastases, closely matching the demographics and disease characteristics of the subjects in our scRNA-seq and bulk RNA-seq analyses (Table S1). This design facilitated identifying cell-specific markers and immune states with high spatial and cellular resolution.

We targeted six key markers to identify major immune cell types and tumor cells, including pan-CK (for tumor cells), CD68 (macrophages), CD56 (NK cells), CD45 (a pan-immune cell marker), CD3E (for T cells), and CD8 (for cytotoxic T cells) using a smaller panel from a larger validated one (Figure 6A).^41,46^ In addition, we selected four markers—CD107a (degranulation marker), PD-1, PD-L1, and ICOS—to capture states of immune activation and cytotoxic potential (Table 5S). This approach allowed us to assess both the localization and activation status of these immune populations, particularly focusing on NK cells and CD8+ T cells. To support our whole slide image analyses (WSI), we employed Cellpose 3.0 for segmentation of the multiplexed immunofluorescence images.^47^ Segmentation objects were categorized into three architectural features—whole slide, intra-tumoral, and peri- tumoral—based on a pathologist’s assessment of the tissue. We then utilized two algorithms as part of our *AstroSuite* package:^42^ 1) *TACIT* (<u>T</u>hreshold-based <u>A</u>ssignment of <u>C</u>ell Types from Multiplexed Imaging Da<u>T</u>a) for cell identification and characterization of cell states, integrating these insights into our analysis of immune cell distribution and activation.^48^ 2)*Astrograph* for data visualization, allowing us to effectively present the spatial patterns and relationships among the different immune cell populations.

**Figure 6.**
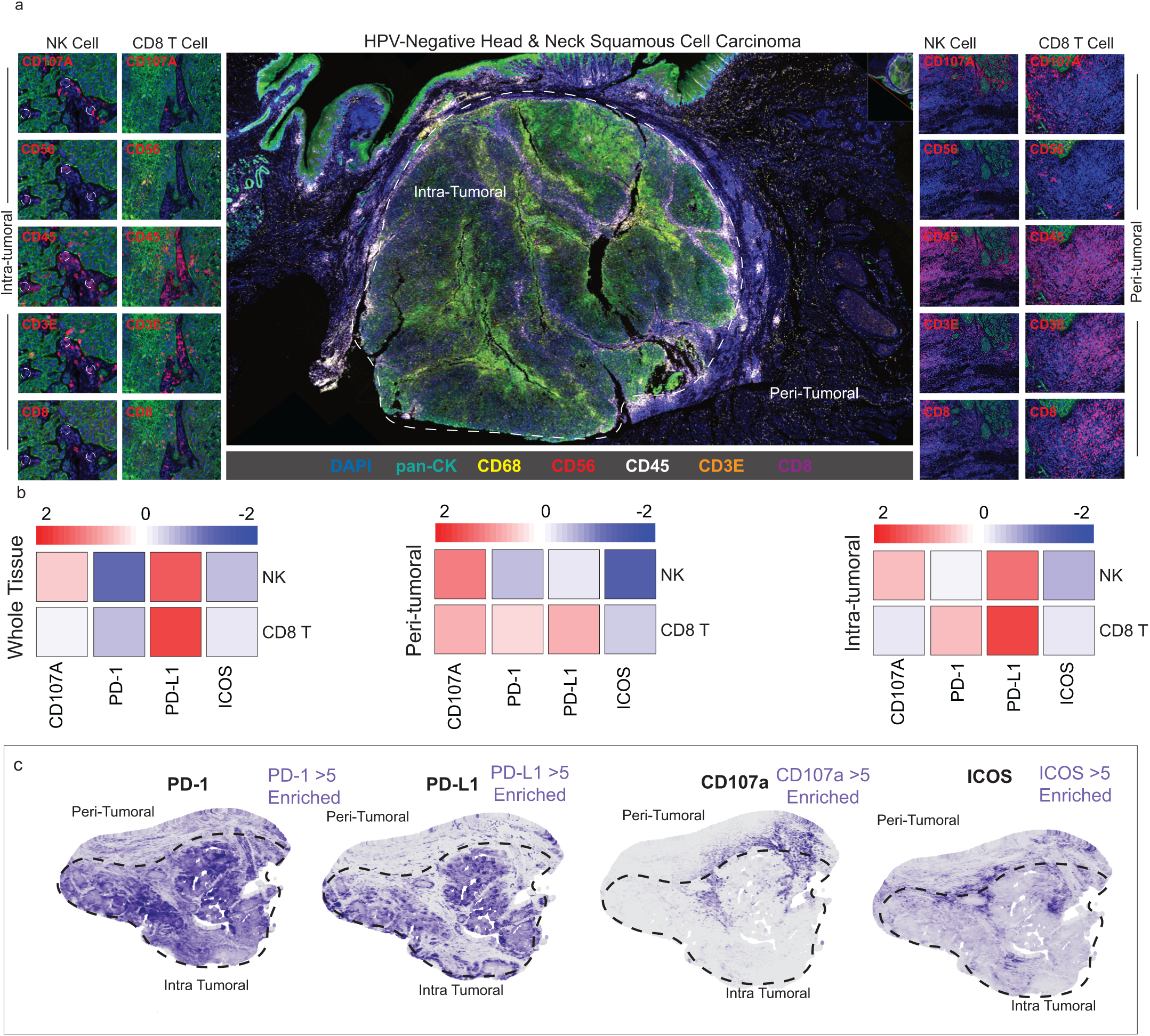
Spatial Proteomics of HPV-Negative HNSCC. The Spatial Multi-IF assay at lower resolution presents a representative area from one of the tumors analyzed in the spatial proteomics study. In this image, intra-tumoral and peri-tumoral regions are highlighted, with analyses focused on compartmentalizing these two subsets within each tumor. The intra- tumoral region (*left)* shows NK cells and CD8+ T cells (*dashed-white circle of central panel*) with lower concentrations of immune markers compared to the peri-tumoral region (*right*) (A). Z-score heatmap representation of immune marker expression levels in NK and CD8+ T cells across different tissue compartments (whole tissue, peri-tumoral, and intra-tumoral regions). (B) Spatial distribution of immune markers in tumor regions. The left panel shows CD107a enrichment, with peri-tumoral areas noted in red and tumor-enriched regions marked in blue. The right panels display PD-1, PD-L1, HLA-A, and ICOS expression maps according to *TACIT* annotation. For each marker, peri-tumoral and intra-tumoral regions are outlined with dashed lines (C). NK – Natural Killer.

In our analysis, we observed notable differences in the spatial distribution and activation states of immune cells both within the tumor core and at the peri-tumoral boundaries. The multi-IF data revealed heterogeneity within the tumor microenvironment, where distinct subsets of NK cells and CD8+ T cells displayed varying levels of cytotoxic and immunosuppressive markers (Figure 6A). In the intra-tumoral regions, a portion of CD8+ T cells exhibited increased expression of PD-1/PD-L1 (Z score = 0.37 and 0.83) compared to peri-tumoral, suggesting an immunosuppressive phenotype that may inhibit cytotoxic activity. In contrast, peri-tumoral areas demonstrated elevated levels of CD107a and ICOS (Figure 6B), indicating active degranulation and potential immune activation among cytotoxic immune cells. These patterns imply a dynamic balance between immune activation and suppression, likely influenced by spatial cues within the tumor microenvironment. The spatial distribution revealed enriched areas of PD-1 and PD-L1 within intra-tumoral regions, while CD107a was predominantly enriched in peri-tumoral areas (Figure 6C). Our findings suggest that the heterogeneity observed across intra-tumoral and peri-tumoral regions drives differential immune cell phenotypes we observed in TICs using scRNAseq.

## Discussion

The TME plays a critical role in cancer progression, therapeutic resistance, and overall patient outcomes. Diverse cell populations within the TME, including immune cells from myeloid and lymphoid lineages, significantly influence tumor behavior through their modulation of the immune response. TICs have been associated with better prognosis and enhanced efficacy of immunotherapy across various cancers, including HNSCC.^49–52^ However, the inherent heterogeneity of the TME, coupled with inter-patient variability and differences between primary tumors and metastatic sites, presents significant challenges in improving treatment outcomes and increasing overall survival rates. Employing advanced techniques such as scRNA-seq and spatial multiomics is essential to gain a comprehensive understanding of tumor heterogeneity and immune dynamics.^53^

Our study builds on the foundation laid by integrated meta-atlases and analyses, including the Human Cell Atlas Initiative/Oral & Craniofacial Bionetwork - Caetano et al. 2022, Easter et al., 2024, and Matuck, Huynh, and Pereira et. al. studies which are critical for understanding the cellular and molecular makeup of human oral and craniofacial tissues in health and disease. Their efforts to map cell type diversity and states using single-cell and spatial multiomics underscore the broader utility of these approaches in biomedical research. The comprehensive atlas developed by the Oral and Craniofacial Bionetwork, which includes factors such as age, sex, and ancestry, aligns with our focus on dissecting immune cell heterogeneity within the TME of HPV-negative HNSCC. Future iterations of these atlases should prioritize the addition of comparisons between healthy and diseased tissues, as observed in our study, to identify novel therapeutic targets and better understand immune dysregulation contributing to tumor progression and metastasis in HNSCC.^41,42,54^

ScRNA-seq technology has increasingly been applied to analyze immune infiltration in the TME of various cancers, including HNSCC.^55^ Notably, Cillo et al. explored the role of CD45+ immune cells in both carcinogen-induced (HPV-negative) and HPV-positive HNSCC. Their study revealed similarities in CD8+ T cell and CD4+ Treg populations between the two subtypes while identifying significant differences in other immune cell types, such as CD4+ T cells, B cells, and myeloid cells. The authors hypothesized that viral antigens in HPV-positive HNSCC may enhance the infiltration of innate immune cells, thereby activating adaptive immune responses, which could explain the observed differences in immune responses and tumor behavior between HPV-positive and HPV-negative HNSCC. Further extending this analysis, Kürten et al. investigated the inflammatory status of the TME in both HPV-positive and HPV-negative HNSCC, uncovering novel fibroblast subsets in the TME of HPV-positive patients. They also emphasized the role of PD-L1+ macrophages in creating an immunosuppressive TME, making these cells critical predictive markers for immunotherapy efficacy. Despite these advances, the specific roles of immune cell subsets during different pathological stages of disease progression and metastasis remain inadequately explored. Our study represents the first integration of scRNA-seq data from PBMCs and TICs in HPV- negative HNSCC, establishing a foundational immunological atlas for this tumor type and evaluating the gene expression of immune cells as they transition from N0 to N+ and T1 to T4 stages.

Our GSEA on the integrated atlases revealed enrichment of the IFN pathway in immune cells of individuals with node stage N+. Notably, no TIC population exhibited high IFN expression (data not shown), indicating TME-dependent modulation and suggesting that high IFN signaling may be primarily expressed by the tumor itself. High IFN signaling can lead to elevated IFI44 expression and immunosuppression, correlating with poorer progression-free survival in HNSCC.^56^ This was accompanied by an increase in cellular interactions predominantly driven by intermediate macrophages and TREM2+ macrophages, which release immunosuppressive signals through IL-10 and TGF-β pathway activation. IL-10 signaling targets dendritic cells (DC2) and intermediate macrophages, both critical antigen-presenting cells (APCs) in activating T lymphocytes for the anti-tumor response. However, IL-10 can inhibit APC activity, facilitating tumor escape and progression.^57,58^ The TGF-β pathway was also positively enriched due to increased expression of its main ligands (TGFB1, TGFB2, TGFB3), likely impacting Tem/Temra cytotoxic T cells via signaling through the TGFBR1-TGFBR2 heterodimer. This enrichment is associated with reduced CD8 T cell immunoscore and a high inflammatory signature, potentially mediated by IFN in HNSCC.^59^ Concurrently, cytotoxic T lymphocytes emerged as primary receptors for signals like IL-2 and CD70, both of which may contribute to the exhaustion phenotype observed in N+ patients. Prior studies have shown that CD70 can induce TIC exhaustion in renal cell carcinoma and TGF-β has been demonstrated to promote CD70 overexpression via the IL-2 pathway in non-Hodgkin lymphomas, leading to an exhausted TIC phenotype characterized by high PD- 1 expression.^60,61^

Based on this data, we hypothesized that N+ tumors had an immune suppressed environment and selected genes associated with NK and cytotoxic T cell activity and modulation to investigate in further detail using pseudotime analysis. We discovered a noticeable increase in the expression of immunomodulatory molecules and a reduction in effector and activating molecules in the tumor, signaling immune suppression. In the NK cell population, we found an increase in the expression of *KLRG1*, a co-inhibitory receptor that regulates NK cell activity, during progression to N+ and *AREG*, a factor which correlates with an unfavorable outcome in the cytotoxic response and is responsible for limiting the expression of Granzymes.^27,62–66^ Consistently, we found reduced *GZMA* expression levels in N+ samples in NK cells. Additionally, the levels of *KIR2DL4*, a receptor which has been described as having a fair prognosis in melanoma, are also reduced in N+ individuals.^67^

Recognizing the significance of the immune environment within the tumor, immunotherapies such as anti-PD1 (Nivolumab) have been approved for treating HNSCC in patients with recurrent or metastatic disease after traditional therapies fail. Nivolumab has demonstrated improved overall survival in individuals with PD-L1 positivity exceeding 1%.^68^ Similarly, Pembrolizumab, another anti-PD1 therapy, has shown improved overall survival compared to chemotherapy with anti-EGFR Cetuximab.^19,20^ Our findings support this, as progression to T4 was marked by increased PDCD1 (PD-1) expression in cytotoxic T lymphocytes. However, the efficacy of these treatments often hinges on the tumor’s PD-L1 expression, with some tumors exhibiting resistance despite PD-L1 positivity.^69^ Thus, exploring alternative therapies is crucial. Greenberg et al. found minimal co-expression of PD-1 and KLRG1 in TICs across various tumor types and demonstrated that administering anti-KLRG1, either alone or with anti-PD1, in murine models of breast, colon, and melanoma led to reductions in tumor volume and metastases.^70^

However, there are currently no clinical trials addressing the use of anti-KLRG1 ABC008 in HNSCC, either alone or in conjunction with anti-PD1. Our data reveal that KLRG1 is upregulated in N+ HNSCC patients, suggesting a mechanism for immune suppression and indicating that metastatic patients could benefit from anti-KLRG1 therapy.

Future investigations should also consider the potential of utilizing two- and three-cell biomarker combinations to enhance the precision of targeted therapies in HNSCC. By identifying specific interactions among immune cell subsets, such as CD8+ T cells, NK cells, and various myeloid cells, we can gain insights into how these cells collaborate or compete in the TME. For example, the presence of a specific combination of immune cell markers could indicate a more effective anti-tumor response or highlight potential resistance mechanisms. This approach not only allows for the identification of more nuanced biomarkers that can predict patient responses to immunotherapy but also facilitates the development of therapies designed to specifically enhance the activity of beneficial immune cells while mitigating the effects of immunosuppressive populations.

Moreover, exploring therapies that simultaneously target both immune and structural cell types within the TME presents a promising avenue for a holistic approach to immunoregulation. By integrating strategies that address the interactions between immune cells and stromal components, such as fibroblasts and extracellular matrix elements, we can create a more favorable environment for immune activation. For instance, therapies that target TGF-β signaling in the stroma may not only reduce immune suppression but also enhance the infiltration and efficacy of cytotoxic T cells and NK cells. Such combinatorial strategies could lead to synergistic effects, improving the overall therapeutic response and reducing tumor progression. Ultimately, a comprehensive understanding of the cellular dynamics within the TME, facilitated by two- and three-cell biomarker profiling, will be instrumental in devising innovative therapeutic approaches that effectively harness the immune system while addressing the structural challenges posed by the tumor microenvironment. In conclusion, embracing this multifaceted strategy may significantly advance the field of immunotherapy and improve outcomes for patients with HPV-negative HNSCC.

## Methods

### Data acquisition and patient characteristics

We integrated two publicly available single-cell RNA sequencing datasets, namely GSE139324 and GSE164690.^21,22^ Clinical metadata and sequencing data were downloaded from the NCBI Gene Expression Omnibus (GEO) repository.^71^ Only HPV negative (HPV-) samples were selected for analysis. Disease staging had been carried out previously in the original publications, following the tumor, node, metastasis (TNM) staging system for HNSCC.^72^ For the dataset GSE139324, we included a total of 17 patients, who were grouped based on tumor staging: T1 (n=1), T2 (n=2), T3 (n=8), and T4 (n=6). Patients with at least one site of lymph node (LN) metastasis were classified as N+ (n=10), whereas patients without LN metastasis were classified as N0 (n=7). For this dataset a total of 34 samples were included for investigation (17 PBMC samples and 17 tissue samples for TICs analysis). Regarding the dataset GSE164690, we included 12 patients who were also allocated to groups based on tumor staging and LN status: T1 (n=1), T2 (n=2), T3 (n=7), and T4 (n=2); N0 (n=7) and N+ (n=5), resulting in a total of 24 samples (12 PBMC samples and 12 tissue samples). All patient information was recovered according to the original publications and is shown in Table S1.^21,22^

### Initial data processing and quality control

The raw FASTQ files for the datasets GSE139324 and GSE164690 were obtained from the NCBI GEO database. The initial transcriptome data were processed using CellRanger version 7.1.0 (10x Genomics),^73^ and sequences were aligned with the human reference genome GRCh38, which FASTA and index files were obtained from the 10x Genomics web page. The output gene-count matrix files were analyzed using the Seurat R package version 5.0.3.^74,75^ Ambient RNA was corrected using SoupX package version 1.6.2 and Doublets were identified using the scDblFinder package version 1.16.0, and a remove.doublet file was generated.^76,77^ The cells in the remove.doublet file were removed from the dataset using Seurat with the command “subset()”. Cells with fewer than 200 genes, with an UMIs above 5 MADs from the median of their sequencing batch, and with more than 10% of mitochondrial RNA genes were excluded.

### Data normalization, dimensionality reduction and clustering

Prior to the clustering and data visualization steps, data were normalized using the NormalizeData function from Seurat, considering default settings, resulting in a log-transformed transcript output per 10,000 reads. The normalized gene/barcode matrix was used to perform dimensionality reduction. Highly variable genes were used to account for the level of variance to perform a principal component analysis (PCA). When performing PCA, we aimed to solely use high quality cells, and therefore sources of sample-specific and cell-specific variation were regressed out. MALAT1, NEAT1, MTRNR, Hemoglobin, ribosomal and mitochondrial genes were removed from the highly variable gene input. PCA was performed using the centered and scaled highly variable gene accordingly by using the “RunPCA” function within Seurat. To visualize the data in a low dimension, uniform manifold approximation and projection (UMAP) was performed, using UMAP-learn python implementation with metric set at “correlation”, for FindNeighbors, nearest neighbor parameter k was set at 10. For Both RunUMAP and findNeighbors functions, the dims parameter was set with several PCs chosen to account for 90% of the variation, 29 for PBMCs and 32 for TICs. Clusters were then identified using the “FindClusters” function from Seurat, the resolution parameter was set at different values to ultimately retrieve a comprehensive clustering based on resolution 2 and visualized on a UMAP plot.

### Cell type identification and annotation

To infer cell types independently to specific clusters, we used CellTypist version 1.6.2, a comprehensive reference database for human immune cells.^23^ Cell types were annotated using the CellTypist model Immune_All_Low version 2, which at the moment of the analysis included 98 different cell types and states from 16 different tissues. This model predicts query cells to be allocated to specific cell type labels using the low-hierarchy classification, which is indicative of a high-resolution prediction. The most probable cell labels were matched with the cell clusters (generated by FindClusters) in our dataset and all other cell type labels from the chosen model were removed. As cell types and states were defined by specific markers, we utilized the ‘Curated markers’ and “Top Model Markers” definitions from CellTypist as a guide to classify each of the cell types and their corresponding possible states, which was followed by the manual selection of the representative markers identified with wilcoxauc() function from presto package version 1.0.0 for each cell type and its corresponding lineage from PBMCs and TICs. Cell types were annotated by joining both datasets and subsequently splitting in TICs and PBMCs, starting from raw counts, but importing to Scanpy the cell clusters, UMAP and PCA embeddings from Seurat analysis, lognormalization in Scanpy was applied before use CellTypist; alternatively, cell annotation of each dataset alone was performed.

### Differential gene expression and gene set enrichment analysis (GSEA)

To identify genes that were differentially expressed between groups, we performed the analysis for each dataset (GSE139324 and GSE164690) using the FindMarkers function in Seurat. Significant differences were identified using the non-parametric Wilcoxon rank sum test. Briefly, observed gene expression values were ranked and the distribution of gene ranks of the N0 group were tested for significance compared to those of the N+ group. Hence, top genes from each cell type in group N0 were compared to the same cell type of group N+. To incorporate solely significant differentially expressed genes (DEGs) in our plot, we set an adjusted p-value threshold at <0.05. To exclude genes with low Log2FC, a default cutoff threshold was set at 0.25 according to Seurat default. All DEGs were then displayed in a jitterplot where those with defined immunological functions among the 20 highest Log2FC in N+ relative to N0 were highlighted. To identify a possible correlation between the functional gene terms in particular groups or clusters and an underlying biological importance of cellular processes, we performed a gene set enrichment analysis (GSEA), that is a computational method that assesses whether a predefined set of genes exhibits statistically significant and concordant differences between biological states, using fgsea R package version 1.28., considering the h.all.v2023.2.Hs.symbols from MSigDB database as reference GMT file.^78^ All genes present in the dataset were ranked using the multiplication product of Log2FC and p-value from the differential gene expression analysis, to get their correlation with the conditions. A threshold was set at an adjusted e-value < 0.05 to include only significantly enriched gene sets.

### Cell-cell communication

To identify intercellular interactions and receptor-ligand signaling networks, we employed the CellChat R package,^43^ a computational tool that infers and visualizes intercellular communication networks from single-cell transcriptomic data. For comparison between N0- and N+ groups, we use the expression data from TICs/PBMCs datasets and generate the analysis for each group per dataset. First, we create a CellChat object from Seurat objects and set “CellChatDB.human” as the database for interactions and reduce the database for keep interactions involved in "Secreted Signaling". Subsequently, to reduce computational costs subsetData() function was used to keep only signaling genes, and overexpressed genes and interactions are identified for being projected over a human protein-protein interaction network. Finally, CellChat computes the communication probability at the signaling pathway level with computeCommunProbPathway() function and calculates the aggregated cell-cell communication network for all cell groups using the aggregateNet() function.

### Gene trajectories

To investigate the pseudotime trajectory ordering of single cells and corresponding genes relevant to each cell type in TICs, we used Monocle 2 v2.30.0,^79,80^ a robust tool designed for single-cell RNA-seq data analysis, that employs a reversed graph embedding (RGE) algorithm to order cells along a pseudotime axis, reflecting their progression through a biological process. Initially, we transform data from Seurat objects, extracting already normalized expression data and metadata for cells and genes. Subsequently, genes that are expressed in at least one cell with a minimum expression of 0.1 (default value) were detected and selected to construct the cell trajectory, using DDRTree method (a RGE algorithm) for dimensionality reduction and orderCells() function for pseudotime values estimation. A newer version, Monocle 3 exists but the authors replaced their representation of pseudotime with a UMAP one that is faster to compute and uses less resource, but it’s more complicated to interpret with our type of data.

### Multiplex Protein Immunofluorescence

Samples for this study were selected from the trial NCT0352942 at the University of North Carolina School of Medicine with input from one trained oral pathologist - BFM. FFPE blocks were cut into sequential 5-micron sections and mounted on SuperFrost Plus slides (Thermo Fisher) for subsequent Phenocycler-Fusion 2.0, H&E, and additional staining. y. All slide preparations used RNA-free water and followed an RNAse-free protocol on the Leica autostainer system. For multiplex immunofluorescence (Multi-IF), we used Akoya Biosciences PhenoCycler Fusion 2.0. Samples were first deparaffinized by a graded ethanol series from 100% to 30%. Antigen retrieval employed AR9 (EDTA) buffer from Akoya Biosciences in a low-pressure cooker for 15 minutes, followed by a 1-hour cooling period. Next, the samples were rehydrated in ethanol for 2 minutes and incubated in staining buffer for 20 minutes. The antibody cocktail buffer was prepared following the Phenocycler Fusion guidelines, with four blockers and nuclease-free water added to the staining buffer. Primary antibodies were diluted at a ratio of 1:200 in this cocktail. The combined antibody solution was applied to slides, which were incubated overnight at 4°C in a Sigma-Aldrich humidity chamber. Post-primary incubation, slides were rinsed in staining buffer for 2 minutes, treated with a post-stain fixative solution (10% PFA in staining buffer) for 10 minutes, and then washed three times in 1X PBS for 2 minutes each. Slides were subsequently immersed in ice-cold methanol for 5 minutes. The final fixative solution (FFS) was prepared as directed in the Phenocycler manual. Slides were incubated with FFS at room temperature for 20 minutes, rinsed with PBS, and transferred to the FCAD machine (Akoya Biosciences) for flow cell mounting. Flow cells were securely affixed to the slides with a 30-second application of high pressure, followed by immersion in PCF buffer for 10 minutes before placement in the Phenocycler Fusion.

Reporters for each antibody were prepared using a Reporter Stock solution combined with 5490 µL of nuclease-free water, 675 µL of 10X PCF buffer, 450 µL of PCF assay reagent, and 27 µL of in-house concentrated DAPI to achieve a 1:1000 DAPI dilution per cycle. This process was applied across two slides at a time, with reporters diluted to 1:50 for each cycle and specific channels. Each report solution was aliquoted (250 µL) into a 96-well plate, sealed with Akoya-provided aluminum foil. Two slides at a time were loaded into the Phenocycler Fusion 2.0 fluidic equipment. Manual area mapping was employed for scanning via the PhenoImager in brightfield mode. Low and high DMSO solutions were prepared as outlined in Index B of the Phenocycler Fusion 2.0 manual.

### AstroSuite

*AstroSuite* features algorithms for spatial biology analysis, including TACIT and Astrograph, previously published tools. TACIT (Threshold-based Assignment of Cell Types from Multiplexed Imaging Data) annotates cell types in spatial omics datasets. This unsupervised algorithm uses a CELLxFEATURE matrix, derived from Cellpose 3.0 segmentation, along with a TYPExMARKER matrix informed by expert marker relevance for cell types. TACIT’s process has two stages: First, cells are grouped into homogeneous Microclusters (MCs) with the Louvain algorithm. Cell Type Relevance (CTR) scores are then calculated, correlating marker intensity with cell type signatures, with higher scores indicating stronger associations. Segmental regression divides CTR scores into relevance clusters, setting a threshold to minimize inconsistent assignments. Cells above this threshold are labeled positive for a cell type. In cases of multiple labels, TACIT uses k-nearest neighbors (k-NN) to resolve ambiguity through deconvolution.

## DATA AVAILABILITY

All data, including links to original raw data from each of the 2 studies can be found at GEO: https://www.ncbi.nlm.nih.gov/geo/. The data can also be analyzed at: https://cellxgene.cziscience.com/collections/065ad318-59fd-4f8c-b4b1-66caa7665409

## CODE AVAILABILITY

Analysis notebooks and CELLxFEATURE matrices for Phenocycler-Fusion 2.0 data are available at: https://github.com/Loci-lab.

## ACKNOWLEDGEMENTS

The results shown here are in whole or part based upon data generated by the TCGA Research Network: https://www.cancer.gov/tcga. Data services in support of the research project were provided by the VCU Massey Comprehensive Cancer Center Bioinformatics Shared Resource. Massey is supported in part with funding from NIH-NCI Cancer Center Support Grant P30CA016059. This work was supported by generous start-up funds from the ADA Science & Research Institute (Volpe Research Scholar Award) and Virginia Commonwealth University to KMB

## CONTRIBUTIONS

KH, SS, JL, KMB and PS conceptualized the project.

RGAG, AR, BFM, KH, CAOBJ, JL, VMC and KMB developed methods for data analysis. RGAG, AR, BFM, CAOBJ and JMMK performed formal analysis.

VMC, KMB and PS acquired financial support.

BFM, NK, SS and KMB supported sample collection.

NK, BFM, KH and KMB performed experimental analysis.

PS and KMB managed and coordinated the research activity planning and execution.

RGAG, AR, JMMK, KMB and PS wrote the original draft.

RGAG, AR, BTR, BFM, JL, KMB and PS reviewed and edited the final manuscript.

## Code Availability

Analysis notebooks and CELLxFEATURE matrices for Phenocycler- Fusion 2.0 data are available at: https://github.com/Loci-lab

---

**Figure S1.** **– Gene expression of marker genes across cell populations**. Bubble plots showing marker genes in X across cell populations in Y in PBMCs (top) and TICs (bottom). The size of the dot represents the percentage of population which expresses the marker gene, and the color represents the average expression from -2 (blue) to +2 (red). PBMC – Peripheral Blood Mononuclear Cells; NK – Natural Killer; pDC – Plasmacytoid Dendritic Cell; Tcm – T Central Memory; Tem – T Effector Memory; Trm – T Resident Memory; Temra – T Effector Memory expressing CD45RA; DC – Dendritic Cell; MAIT – Mucosal Associated Invariant T Cell; TIC – Tumor Infiltrating Cells.

**Figure S2.** **- Clustering and proportions of immune cell populations in PBMCs from GSE139324 and GSE164690**. UMAPs stained by Cell Types (A) and Pathological Node (B) in PBMCs. Bar graphs show proportions of cell types across samples (C) or Pathological Node (D) from GSE139324. UMAPs stained by Cell Types (E) and Pathological Node (F) in PBMCs. Bar graphs show proportions of cell types across samples (G) or Pathological Node (H) from GSE164690. HNSCC – Head and Neck Squamous Cell Carcinoma; PBMC – Peripheral Blood Mononuclear Cells; NK – Natural Killer; pDC – Plasmacytoid Dendritic Cell; Tcm – T Central Memory; Tem – T Effector Memory; Trm – T Resident Memory; Temra – T Effector Memory expressing CD45RA; DC – Dendritic Cell; MAIT – Mucosal Associated Invariant T Cell; UMAP – Uniform Manifold Approximation and Projection;

**Figure S3.** - Clustering and proportions of immune cell populations in TICs from GSE139324 and GSE164690. UMAPs stained by Cell Types (A) and Pathological Node (B) in TICs. Bar graphs show proportions of cell types across samples (C) or Pathological Node (D) from GSE139324. UMAPs stained by Cell Types (E) and Pathological Node (F) in PBMCs. Bar graphs show proportions of cell types across samples (G) or Pathological Node (H) from GSE164690. HNSCC – Head and Neck Squamous Cell Carcinoma; NK – Natural Killer; pDC – Plasmacytoid Dendritic Cell; Tcm – T Central Memory; Tem – T Effector Memory; Trm – T Resident Memory; Temra – T Effector Memory expressing CD45RA; DC – Dendritic Cell; MAIT – Mucosal Associated Invariant T Cell; UMAP – Uniform Manifold Approximation and Projection; TIC – Tumor Infiltrating Cells.

**Figure S4.** **– Proportion of cells between N status and Pathological stages across cell populations**. Box and whiskers with jitterplots representing cell populations proportions between N status (A) or Pathological Stages (B) from PBMCs, and between N status (C) or Pathological Stages (D) from TICs. Student’s t Test. NK – Natural Killer; pDC – Plasmacytoid Dendritic Cell; Tcm – T Central Memory; Tem – T Effector Memory; Trm – T Resident Memory; Temra – T Effector Memory expressing CD45RA; DC – Dendritic Cell; MAIT – Mucosal Associated Invariant T Cell.

**Figure S5.** **- Differentially expressed genes at the single cell level in PBMCs**. Jitterplots indicating differentially expressed genes between N status (A) and Pathological Stages (B). The size of each dot represents the percentage of cells expressing the gene. Cell types have been highlighted by color. Genes were ranked based on the average Log2FC. Highlighted genes are those that showed the highest differential expression with known immunological function among the 20 genes with highest Log2FC in each cell annotation. The black line represents Log2FC of zero. The dotted line represents the Log2HR threshold of 0.25. HNSCC – Head and Neck Squamous Cell Carcinoma; PBMC – Peripheral Blood Mononuclear Cells; NK – Natural Killer; pDC – Plasmacytoid Dendritic Cell; Tcm – T Central Memory; Tem – T Effector Memory; Trm – T Resident Memory; Temra – T Effector Memory expressing CD45RA; DC – Dendritic Cell; MAIT – Mucosal Associated Invariant T Cell.

**Figure S6.** Bulk RNAseq data are not able to represent these findings. - Graphs showing disease-free survival in Y according to Status N (A) or Pathological (B). Gene expression of genes marked by Status N (A) or Pathological (B). Data deposited from the TCGA-HNSC project. Ln – Lymph Node.

**Table S1.** Patient demographic and tumor data. . Patient demographic and tumor information for single cell, bulk and spatial samples.

**Table S2.** Differential gene expression for TICs and PBMCs based on node status and pathological stage. Average log2 fold change and p values for differentially expressed genes across cell types. Analysis reported for T3/T4 vs T1/T2 and N+ vs N0 sample across both TICs and PBMCs.

**Table S3.** GSEA for TICs and PBMCs based on node status and pathological stage. . Biological pathways and corresponding p-value/ adjusted values for Gene Set Enrichment Analysis across cell types. Analysis reported based on pathological stage and node status across both TICs and PBMCs.

**Table S4.** Number of outgoing/ incoming signals received based on node status. . Number of outgoing signals from intermediate and TREM 2+ macrophages and number of incoming signals for Temra/Tem Cytotoxic T Cells and Trm/Tem Cytotoxic T Cells based on node status.

**Table S5.** Individual cell types were identified using the TACIT annotation tool. Marker intensities, quantified through immunofluorescence, were normalized using Z-scores, enabling a comparative assessment of cytotoxic and immunosuppressive features across different cell populations. The analysis uncovered considerable diversity, with distinct subgroups of NK and CD8+ T cells demonstrating a wide range of functional marker profiles.

