## Supplementary figures and images for "The Single-Cell Landscape of Peripheral and Tumor-infiltrating Immune Cells in HPV- HNSCC"

### Supplemental Figure 1

#### Supplemental Figure 1

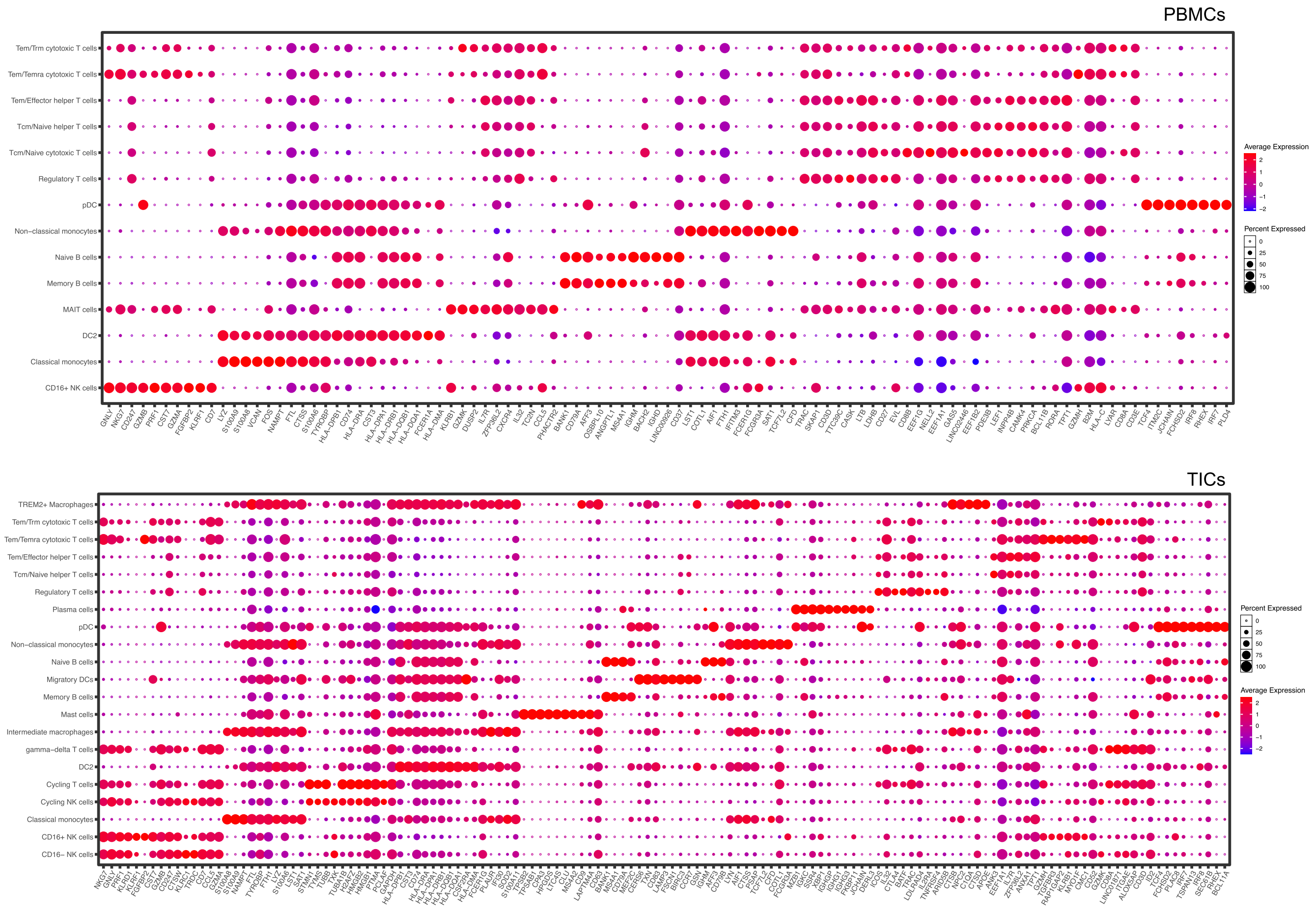

### Supplemental Figure 2

#### Supplemental Figure 2

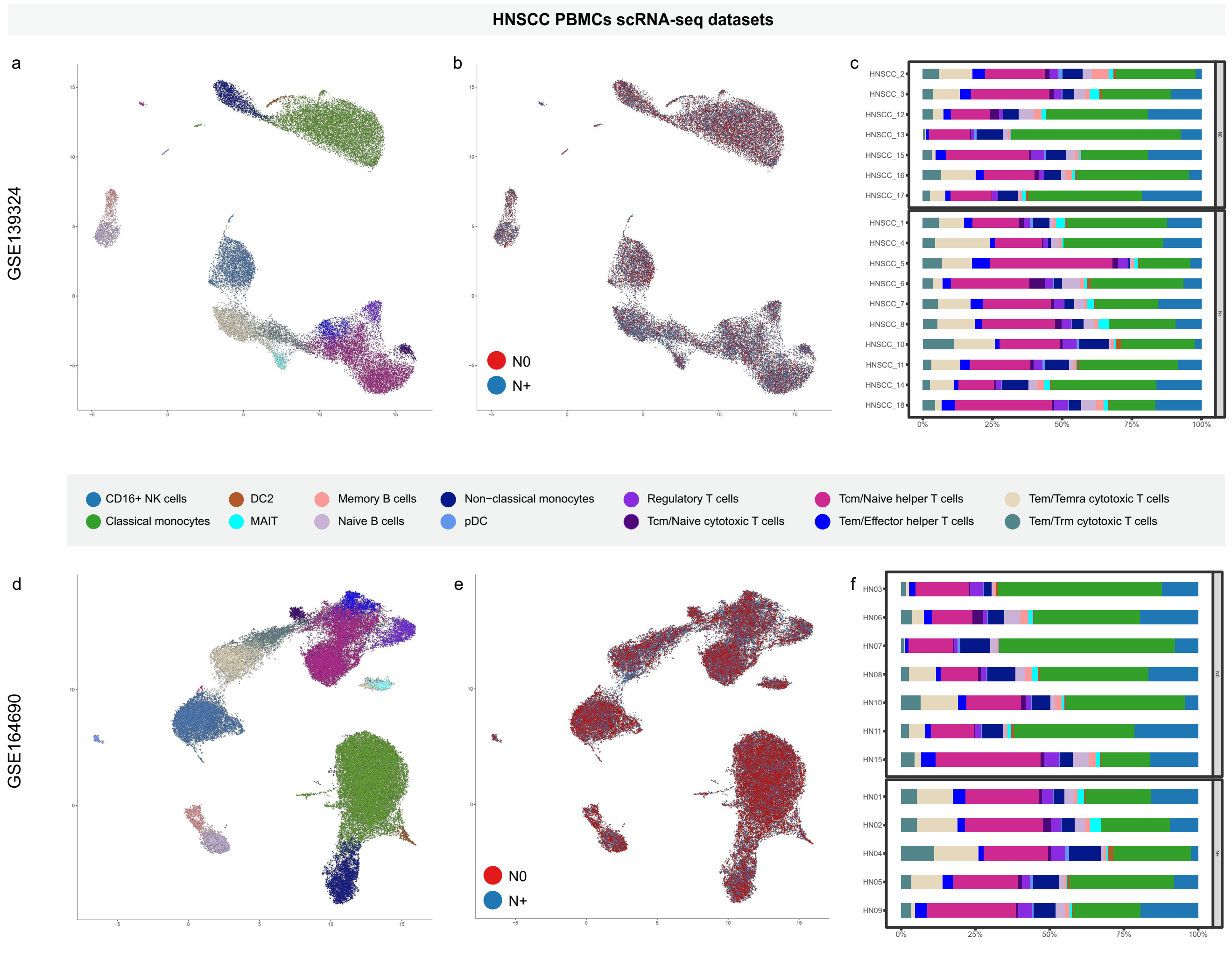

### Supplemental Figure 3

#### Supplemental Figure 3

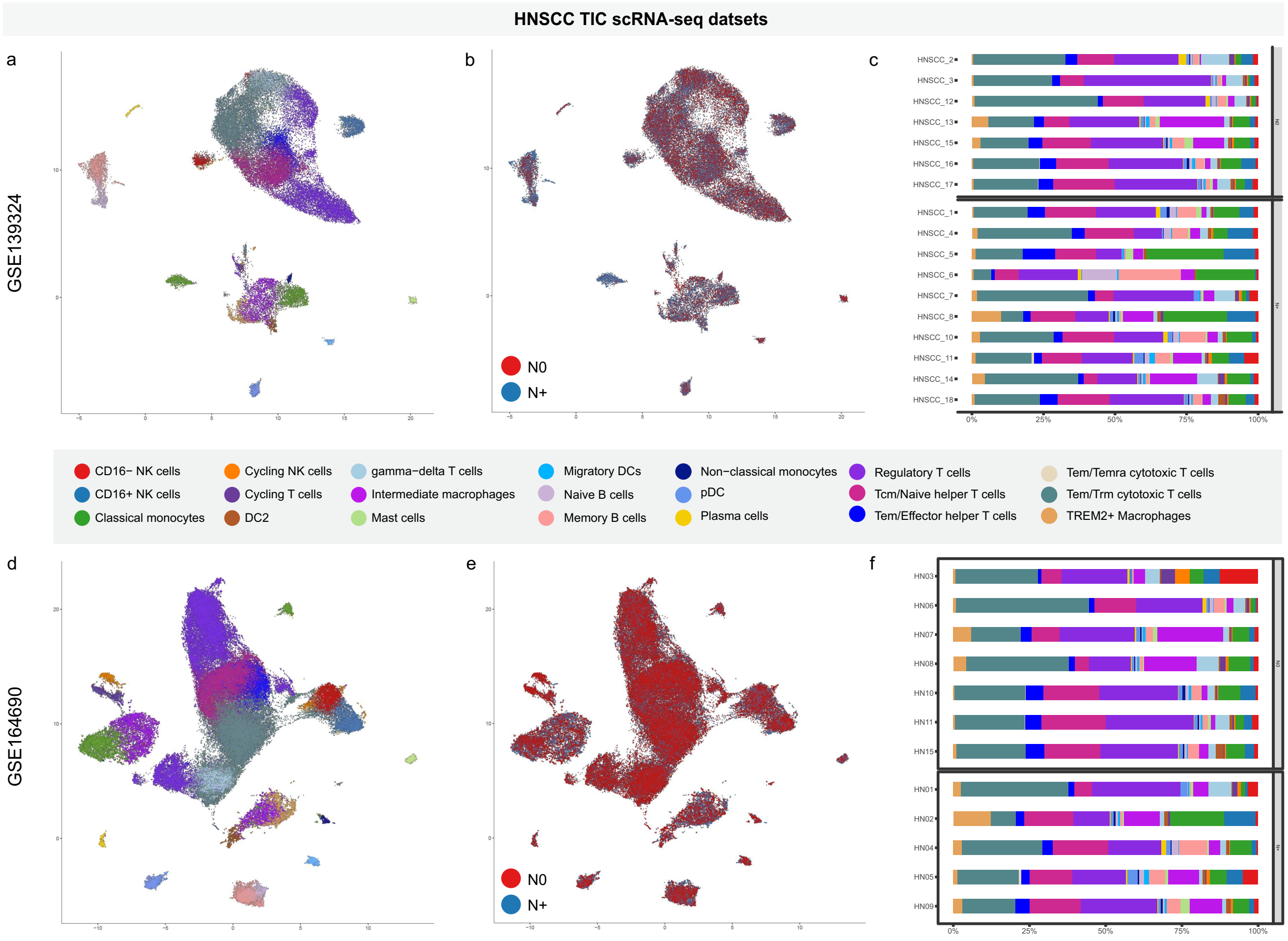

### Supplemental Figure 4

Supplemental Figure 4

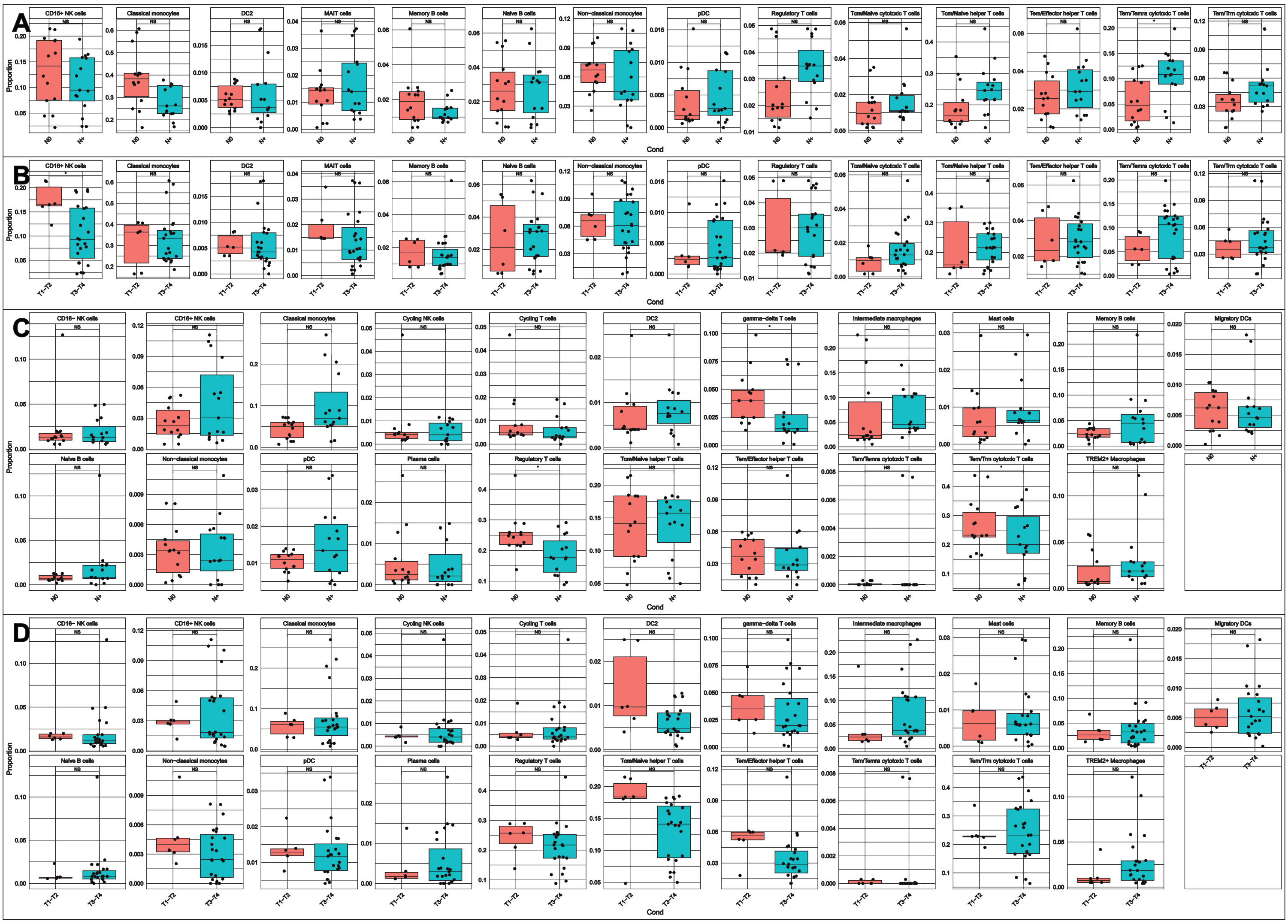

### Supplemental Figure 5

#### Supplemental Figure 5

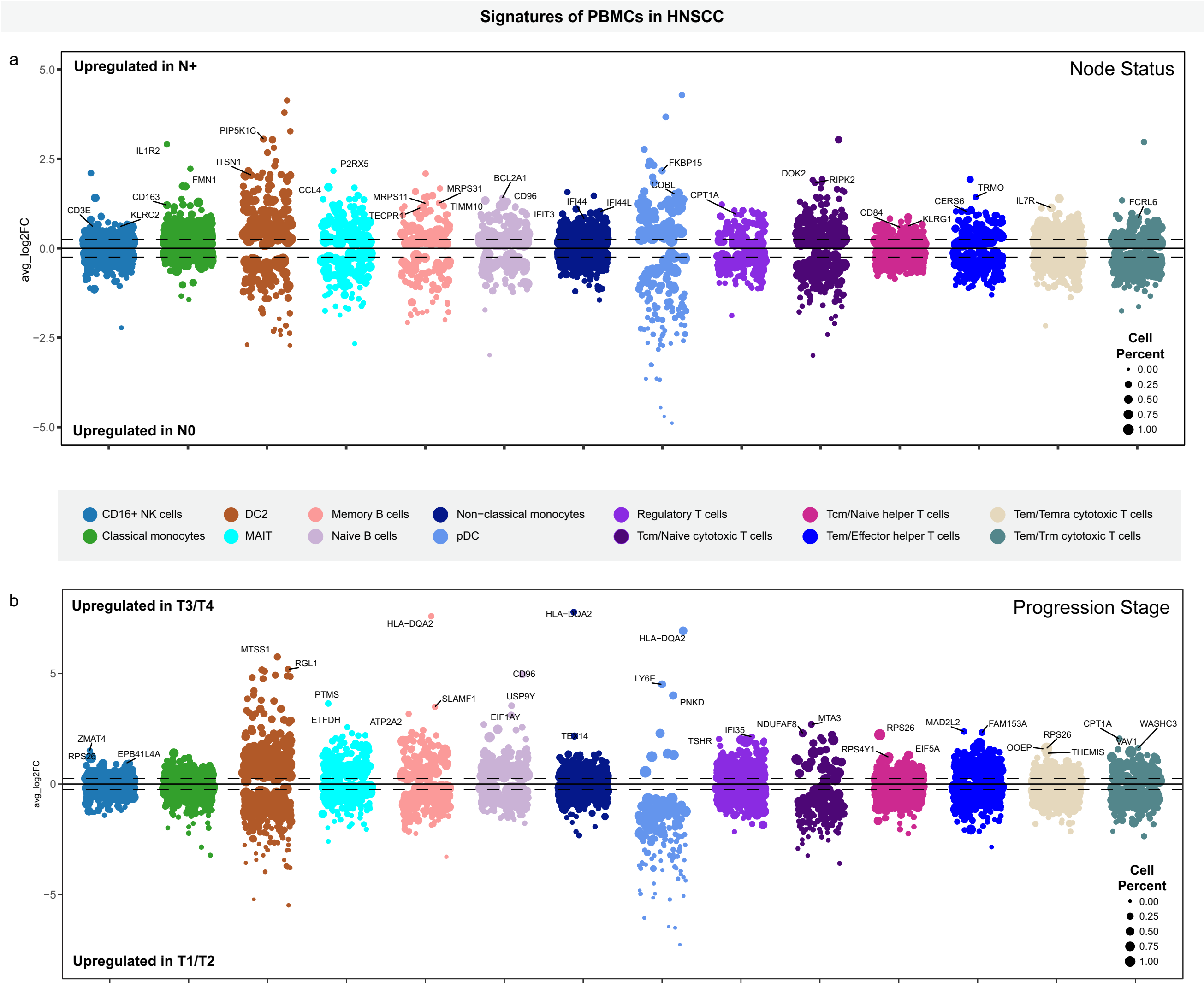

### Supplemental Figure 6

Supplemental Figure 6

a

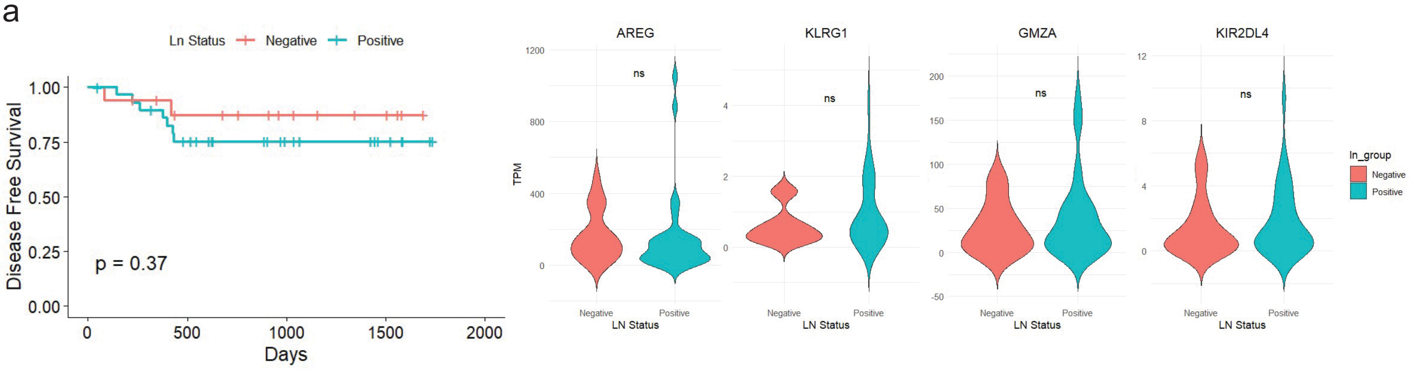


b

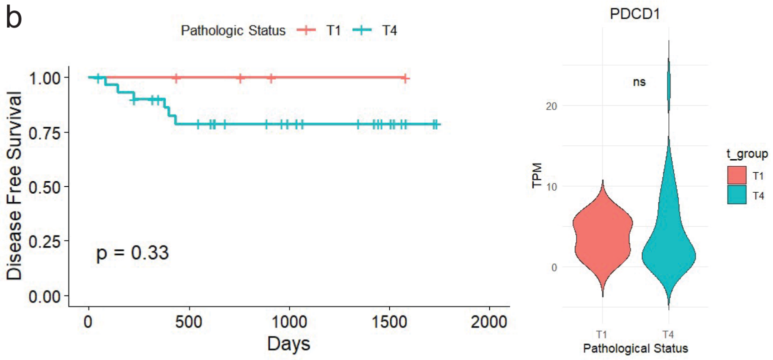
